# A multi-transcriptomics approach reveals the coordinated action of the endoribonuclease DNE1 and the decapping machinery in orchestrating mRNA decay

**DOI:** 10.1101/2024.01.31.578142

**Authors:** Aude Pouclet, David Pflieger, Rémy Merret, Marie-Christine Carpentier, Marlene Schiaffini, Hélène Zuber, Dominique Gagliardi, Damien Garcia

## Abstract

Decapping is a crucial step of mRNA degradation in eucaryotes and requires the formation of the holoenzyme complex between the decapping enzyme DCP2 and the decapping enhancer DCP1. In Arabidopsis, we recently identified DNE1, a NYN domain endoribonuclease, as a direct protein partner of DCP1. The function of both DNE1 and decapping are necessary to maintain phyllotaxis, the regularity of organ emergence in the apex. In this study we combined *in vivo* mRNA editing, RNA degradome, transcriptomics and small RNA-omics to identify targets of DNE1 and study how DNE1 and DCP2 cooperate in controlling mRNA fate. Our data reveal that DNE1 mainly contacts and cleaves mRNAs in the CDS and has sequence cleavage preferences. We found that DNE1 targets are also degraded through decapping, and that both RNA degradation pathways influence the production of mRNA-derived siRNAs. Finally, we detected mRNA features enriched in DNE1 targets including RNA G-quadruplexes and translated upstream-ORFs. Combining these four complementary high-throughput sequencing strategies greatly expands the range of DNE1 targets and allowed us to build a conceptual framework describing the influence of DNE1 and decapping on mRNA fate. These data will be crucial to unveil the specificity of DNE1 action and understand its importance for developmental patterning.

## Introduction

Eucaryotic cells possess a large panel of general and specific mRNA degradation activities to precisely set mRNA homeostasis and fine tune gene expression programs. These activities include: the mRNA decapping complex formed by the enzyme Decapping 2 (DCP2) and decapping activators including Decapping 1 (DCP1) and Enhancer of decapping 4 (EDC4) (He and Jacobson, 2022; Vidya and Duchaine, 2022); 5’-3’ and 3’-5’ exoribonucleases including the exoribonuclease XRN1 and the RNA exosome complexes (Schmid and Jensen, 2019; Krempl et al., 2023); several endoribonucleases including ARGONAUTE proteins involved in RNA silencing (Poulsen et al., 2013), SMG6 involved in nonsense-mediated decay and MARF1 a NYN domain endoribonuclease which acts together with proteins involved in decapping to regulate the degradation of specific transcripts (Nishimura et al., 2018; Boehm et al., 2021). DCP2 and exoribonucleases are general factors involved in bulk mRNA degradation but are also involved in mRNA quality control and regulatory pathways such as nonsense-mediated decay (NMD) or miRNA-mediated gene silencing (Rehwinkel et al., 2005; He and Jacobson, 2022). In plant*s*, most of the activities cited before exist including the decapping enzyme DCP2 in association with the decapping activators DCP1, VARICOSE (VCS) and EXORIBONUCLEASE 4 (XRN4), the plant homologues of EDC4 and XRN1, respectively, and the plant 3’-5’ RNA exosome (Souret et al., 2004; Zhang et al., 2015; Lange and Gagliardi, 2022). A specificity of plant is the tight link between RNA degradation and RNA silencing. This phenomenon is due to the use in plants of a dedicated RNA silencing amplification machinery to fight against viruses and other invading elements like transposons (Lopez-Gomollon and Baulcombe, 2022). A key challenge inherent to RNA silencing amplification is to avoid targeting of its own mRNAs by this defense mechanism. RNA degradation activities carried by DCP2, XRN4, as well as the RNA exosome, protect the transcriptome against RNA silencing activation in plants. Indeed, several mutations in RNA degradation factors lead to the production of mRNA-derived siRNAs, often resulting in developmental defects (Gregory et al., 2008; De Alba et al., 2015; Branscheid et al., 2015; Lam et al., 2015; Zhang et al., 2015; Lange et al., 2019).

In the model plant *Arabidopsis thaliana*, we recently identified DNE1 an endoribonuclease associated with the decapping enhancers DCP1 and VCS and co-purifying with the RNA helicase UPF1 required for NMD. DNE1 is the closest homologue of MARF1 and is composed of a NYN endoribonuclease domain associated with two OST-HTH domains predicted as RNA binding modules. We found that DNE1 together with decapping are crucial for the precise developmental patterns appearing during flower emergence in the shoot apex, a phenomenon called phyllotaxis (Schiaffini et al., 2022). A recent degradome analysis by genome-wide mapping of uncapped and cleaved transcripts (GMUCT; (Willmann et al., 2014; Carpentier et al., 2021)) identified 224 mRNAs producing DNE1-dependent RNA degradation intermediates (Nagarajan et al., 2023). A main achievement of this study was the identification of the first set of mRNAs targeted by DNE1. Yet, the full spectrum of DNE1 mRNA targets remains to be discovered, as well as the interplay between DNE1 and other RNA degradation pathways. In the present study we combined four complementary high throughput sequencing strategies to identify mRNAs directly bound and processed by DNE1 and to understand how this endoribonuclease coordinates its action with the decapping enzyme DCP2 to orchestrate mRNA decay.

First, to identify mRNAs directly in contact with DNE1, we used HyperTRIBE, an *in vivo* RNA editing method in which DNE1 was fused to the catalytic domain of the adenosine deaminase ADAR (Rahman et al., 2018; Arribas-Hernández et al., 2021). In order to define which of these mRNAs were processed by DNE1 we applied a second and complementary approach and analyzed the mRNA degradation patterns influenced by DNE1 using GMUCT. For this approach, we adapted an existing bioinformatic pipeline for normalization and statistical analysis of GMUCT datasets. Using this pipeline, we compared GMUCT datasets for *xrn4* and *xrn4 dne1* mutants and identified more than 1200 loci for which 5’ monophosphate mRNA fragments (5’P) are produced in a DNE1-dependent manner. This result indicates that DNE1 targets a larger repertoire of mRNAs than previously described. In addition, we also identified that DNE1 limits the accumulation of decapped RNA degradation intermediates of some of its targets indicating dual targeting and coordinated action of DNE1 and decapping. To study this coordinated action of DNE1 and decapping, we analyzed mutants affected in both DNE1 and DCP2 using transcriptomics and small RNA-omics approaches. Our results indicate that the cooperation of DNE1 and DCP2 influences the steady state level of several mRNAs and the production of mRNA-derived siRNAs. Overall, our multi-transcriptomics strategy provides an extended list of DNE1 targets, identified several mRNA features enriched in DNE1 targets and identifies nucleotide preferences for DNE1 cleavage. We provide evidences of the redundancy between the action of DNE1 and decapping in controlling mRNA fate and in protecting mRNAs against RNA silencing activation. Finally, we propose a model of the coordinated action of DNE1 and decapping as a conceptual framework, an important step towards the understanding of how DNE1 and DCP2 cooperate in the regulation of gene expression and in the control of faithful developmental patterns in the shoot apex.

## Results

### Identification of mRNAs associated with DNE1 by mRNA *in vivo* editing

In order to identify mRNAs in direct contact with DNE1, we used the *in vivo* RNA editing strategy HyperTRIBE (Fig.1; Rahman et al., 2018; Arribas-Hernández et al., 2021). For this purpose, we generated Arabidopsis transgenic lines expressing the catalytic domain of the adenosine deaminase ADAR from *Drosophila melanogaster* (thereafter called ADAR) fused to either WT DNE1 or to a DNE1 catalytic mutant (DNE1^D153N^; Fig. 1A). The rationale for the use of the catalytic mutant DNE1^D153N^ was to improve the efficiency of mRNA target edition by limiting their degradation by DNE1 and by increasing the dwelling time of DNE1 on its targets. For this experiment five independent transgenic lines of each construct, considered as five biological replicates were analyzed by RNA-seq and compared with plants expressing an unfused version of ADAR, used as a control as previously described (Arribas-Hernández et al., 2021). This analysis resulted in the identification of 322 and 2268 edited mRNAs by DNE1 and DNE1^D153N^ respectively (Fig. 1B, Supplemental Data Set S1). As expected, most mRNAs (306/322) identified using ADAR-DNE1 were also present in the ADAR-DNE1^D153N^ dataset. The catalytic mutant led to a higher editing efficiency than the WT, in agreement with our initial hypothesis. Strikingly, more than 80% of the editions by DNE1 occurred within CDS with both DNE1 and DNE1^D153N^ (86.1% and 83.4% respectively, Fig. 1C). This result suggests that DNE1 interacts mainly with transcripts internally and not at the 5’ extremity as could be anticipated from its interaction with decapping activators. This preferential internal contact with mRNAs can be visualized on selected transcripts (Fig. 1D, Supplemental Fig. S1). Theoretically, we can envision two alternative scenarios for mRNAs contacting DNE1, either they are in contact with DNE1 and cleaved, or they are in contact with DNE1 but not cleaved.

**Figure 1.**
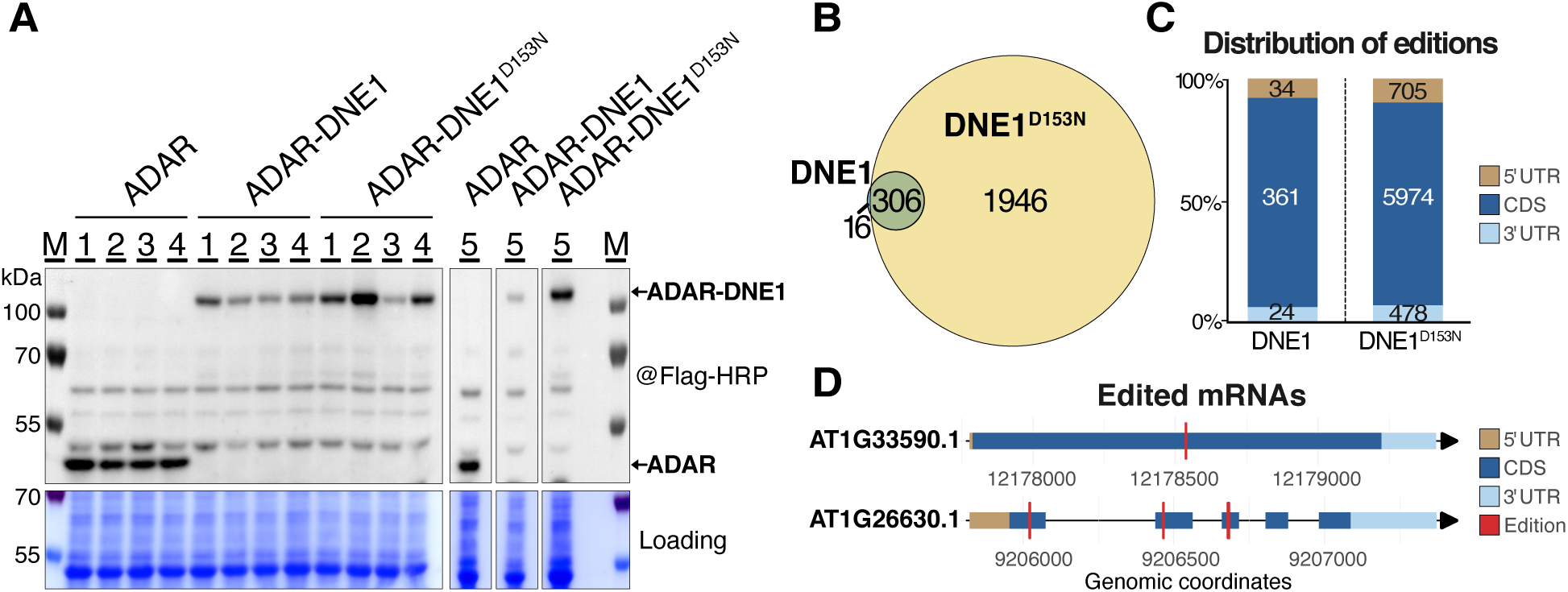
*In vivo* editing using HyperTRIBE identifies mRNA in direct contact with DNE1. (A) Western blot showing the protein accumulation in transgenic lines used for HyperTRIBE and expressing either the ADAR catalytic domain (ADAR) used as a control or protein fusions between DNE1 and ADAR. (B) Venn diagram showing the overlap in loci edited by ADAR-DNE1 or ADAR-DNE1^D153N^. Significant A to G editions were considered with adjpv<0.01, Log2FC>1 and a minimum of 10 reads. (C) Distribution of editions by DNE1 and DNE1^D153N^ on mRNAs. (D) Schemes showing the editions by ADAR-DNE1^D153N^ on two transcripts (additional examples are shown in Supplemental Fig. S1).

### Analysis of mRNA degradation patterns upon DNE1 inactivation implies a dual targeting by DNE1 and decapping

To discriminate between these two scenarios, and gain further insights on the mode of action and targets of DNE1, we performed degradome analysis using GMUCT (Fig. 2). Our experimental setup allows the use of efficient methods for normalization and statistical analysis for target discovery and to quantify all RNA fragments, including the most abundant and secondary 5’P giving access to the complete DNE1 dependent RNA degradation patterns. Differential RNA degradation patterns were identified by adapting the DEXseq method, originally developed to analyze differential splicing patterns (Anders et al., 2012), to analyze GMUCT datasets obtained from biological triplicates. In this analysis, we considered every 5’P identified for a given transcript and compared these fragments between two genetic conditions. The analysis was performed comparing *xrn4* to *xrn4 dne1* in order to work in backgrounds in which 5’P, including those arising from DNE1 activity as an endoribonuclease, are stabilized and increase the probability to detect them using GMUCT. We filtered low covered 5’P by removing positions where the mean RPM of the 3 biological replicates is lower than 1 RPM in all conditions. After differential analysis using DEXSeq, we kept positions with a Log2FC≥1 or Log2FC≤1 and adjusted p-value (adjPv) <0.05. Using this method, we identified 1475 transcripts with differential degradome patterns in *dne1 xrn4* (Fig. 2A, 2B, 2C, Supplemental Data Set S2). The main pattern observed was downregulation of 2631 fragments arising from 1296 individual loci upon mutation of DNE1 in *xrn4* background. This observation implies that some loci accumulate several DNE1 dependent fragments. These fragments are expected to include both direct DNE1 cleavage products and the most stable mRNA degradation intermediates arising from these fragments. This result supports the previous conclusion that DNE1 acts as a *bona fide* endoribonuclease targeting mRNAs, leading to the production of RNA degradation products with 5’-P extremities (Nagarajan et al., 2023). As previous work identified 224 loci producing DNE1-dependent 5’P RNA degradation intermediates with GMUCT, our experimental setup and bioanalysis pipeline greatly expand the spectrum of putative direct DNE1 targets. Examples of these downregulated RNA fragments can be visualized along the transcripts (Fig. 2B, Supplemental Fig. S2A). One particularity of our analysis is to identify significantly downregulated 5’P including both the main RNA degradation intermediate and secondary RNA fragments. Interestingly, 50% of the loci identified previously (111/224; Nagarajan et al., 2023) are present in our dataset validating the efficiency of our method to identify DNE1 targets. To have a global view of the position of these DNE1 dependent RNA degradation patterns, we determined their distribution and compared with the overall accumulation of 5’P. We found that the proportion of downregulated fragments was increased in CDS and 3’UTR compared to all fragments (Fig. 2D), which supports cleavage by DNE1 mostly in the CDS but also in 3’UTR.

**Figure 2.**
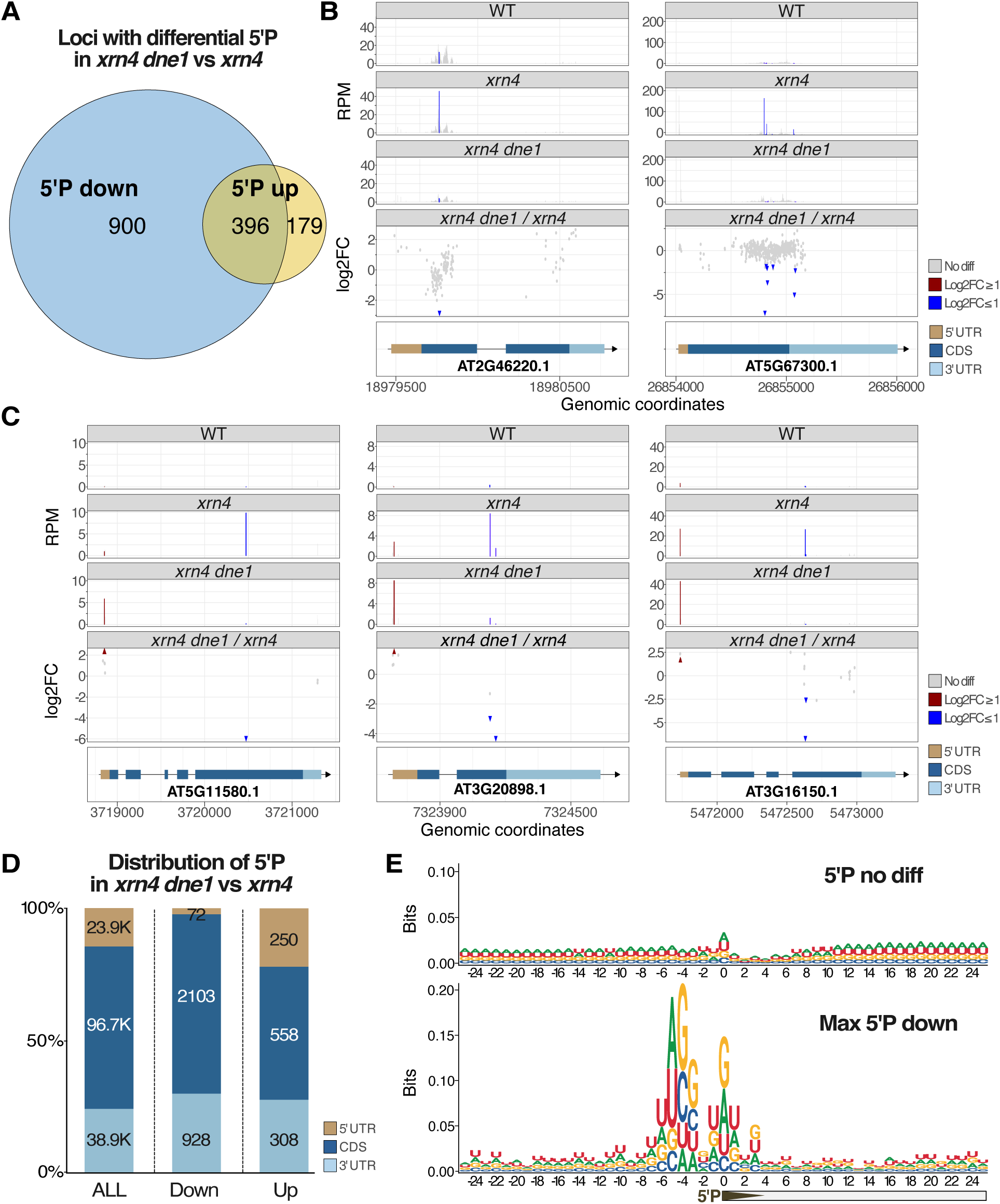
Degradome analysis by GMUCT identifies two opposite trends on DNE1 targets upon mutation in DNE1. (A) Venn diagram showing the output of a differential GMUCT analysis between *dne1 xrn4* and *xrn4* and displaying the overlap between loci showing upregulated and downregulated 5’P fragments. (B) Plots showing the repartition of downregulated 5’P on two loci presenting only downregulated 5’P in *dne1 xrn4*. (C) Plots showing the repartition of 5’P on three loci presenting both downregulated and upregulated 5’P in *dne1 xrn4*. Differential 5’P were considered with Log2FC≥1 or Log2FC≤1 and Pv<0.05 following the DEXseq analysis. Datasets from the three biological replicates were pooled to generate the graphs presented in B and C. (D) Histogram showing the distribution on mRNAs of 5’P depending on their behavior in *dne1 xrn4*. (E) Analysis of the nucleotide composition around the 1295 main DNE1 dependent 5’P site using a sequence logo. The upper panel shows a control sequence logo produced using unchanged 5’P sites in *dne1 xrn4* coming from the 1295 loci producing DNE1 dependent 5’P. The lower panel shows the same analysis using the main DNE1 dependent 5’P from each locus. Position 0 represents the first nucleotide of the 5’P as sequenced in GMUCT.

Somewhat counterintuitively, we also found that 575 transcripts showed increased 5‘P when DNE1 is mutated. Interestingly almost 70% of these transcripts (396/575) were also showing decreased RNA fragments with 5’ end at distinct positions on the transcript (Fig. 2A). Such dual up and down patterns can be visualized along the transcripts (Fig. 2C, Supplemental Fig. S2B). When we compared the localization of upregulated versus downregulated 5’P along transcripts, we observed that the proportion of upregulated 5’P is ten times more important in the 5’UTR than the downregulated 5’P (Fig. 2D). This difference suggests that upregulated 5’P are more prone to occur close to the TSS, some of them could represent decapped fragments or be secondary fragments produced from decapped fragments. To test this hypothesis, we looked in our GMUCT data for fragments identified as decapped sites by C-PARE (Nagarajan et al., 2019). Among our 155 100 GMUCT sites, 14 384 were identified as decapped sites in C-PARE. Most of these sites (14 247) do not change upon mutation of DNE1, indicating that DNE1 does not globally influence decapping. Interestingly, 137 of these sites change when DNE1 is mutated with a predominance of upregulated (124) versus downregulated (13) sites. Therefore, mutation in DNE1 can lead to an increased accumulation of decapping intermediates. Upregulated 5’P occur mainly (70%) on transcripts showing downregulated 5’P at other location, indicating the dual targeting by DNE1 and decapping. This trend can be visualized on many loci including AT5G11580, AT3G20898 and AT3G16150 for example (Fig. 2C, Supplemental Fig. S2B). As we analyzed the complete RNA degradation patterns including main and secondary sites, some upregulated 5’P likely represent secondary 5’P arising from degradation of decapped intermediates. Such examples can be visualized on transcripts presenting many 5’P like AT1G22190 for example (Supplemental Fig. S2B). The RNA degradation patterns with 5’P accumulating more in *xrn4 dne1* generaly occur upstream of decreased 5’P fitting the idea that upregulated fragments derive from decapped mRNAs and downregulated fragments derived from DNE1 endoribonucleolytic cleavage either in CDS or 3’UTR. In conclusion, our experimental setup and exhaustive analysis of DNE1-dependent RNA degradation patterns greatly expand the spectrum of putative DNE1 targets and highlights the coordination of the action of DNE1 and decapping.

### Biased nucleotide composition at DNE1 cleavage sites suggests sequence cleavage preferences

To investigate a potential sequence cleavage preference for DNE1, we analyzed the nucleotide composition in the vicinity of the main DNE1 dependent fragments. A nucleotide logo was produced 25 nt before and after the 5’ extremity of these fragments on the 1296 loci with downregulated 5’P in GMUCT. Interestingly, whereas no bias is observed in a control analysis performed on DNE1-independent 5’P, a significant deviation from a random nucleotide composition appears in the close vicinity of these 1296 cleavage sites. The nucleotide bias observed for downregulated 5’P clearly appears both before and after the 5’P extremity at positions −3 to −6 and −1 to 1 (Fig. 2E). The most extreme values appear at nucleotides −4, −3 and 0 with 46.7, 44.2 % and 38.6% of G respectively, a strong deviation from the 25,4% of G observed when considering the whole region. This non-random sequence composition strongly suggests a sequence preference for DNE1 cleavage activity.

### Analysis of HyperTRIBE and GMUCT data identifies mRNA features enriched in DNE1 targets

We then compared the data obtained by HyperTRIBE with data obtained by GMUCT (Fig. 3). We found that *ca* 22% of the transcripts identified as DNE1 targets by GMUCT (those producing less 5’P fragments in *xrn4 dne1*) were edited by DNE1-D153N (288/1296) identifying them as in direct contact and processed by DNE1 (Fig. 3A). We investigated the presence of specific features in mRNAs identified in these two approaches. Because G-rich motifs were previously identified in DNE1 targets (Nagarajan et al., 2023) and because the first OST-HTH domain of DNE1 was found to interact with G-rich and RNA G-quadruplex structures (rG4) in vitro (Ding et al., 2020), we first looked for the overlap between HyperTRIBE and experimentally validated loci containing rG4 (Yang et al., 2020); Fig. 3B). Interestingly we found that 516 mRNAs directly in contact with DNE1 in HyperTRIBE were containing experimentally validated rG4 in rG4-seq. To determine if DNE1 targets identified by HyperTRIBE and GMUCT were enriched for specific features, we looked at the distribution of diverse mRNA features among these loci, including CDS, UTR length and intron number (Fig. 3C). Whereas no consistent changes were observed between the different lists for CDS and intron numbers, DNE1 targets identified by these methods seemed to systematically harbor slightly longer UTRs. Because of these longer UTRs and the presence of mRNA with rG4 among DNE1 targets, we tested whether the proportion of transcripts containing translated uORFs in 5’UTR (Ribo-seq data from (Hu et al., 2016)) or validated rG4 (rG4-seq data from (Yang et al., 2020)) was higher among DNE1 targets compared to all transcripts expressed in similar tissues either seedlings or flowers. Interestingly, we observed a significantly higher proportion of mRNA containing translated uORFs and rG4 among identified DNE1 targets (Fig. 3D). Strikingly, for both the strongest enrichments were observed for DNE1 targets identified in common between GMUCT and HyperTRIBE, reinforcing the relevance of these features (Fig. 3C, 3D, Supplemental Data Set S6 and S7). Overall, this comparison identifies a set of 288 transcripts directly in contact and processed by DNE1 and reveals that these targets of DNE1 validated by two independent techniques, are enriched in rG4 and translated uORFs.

**Figure 3.**
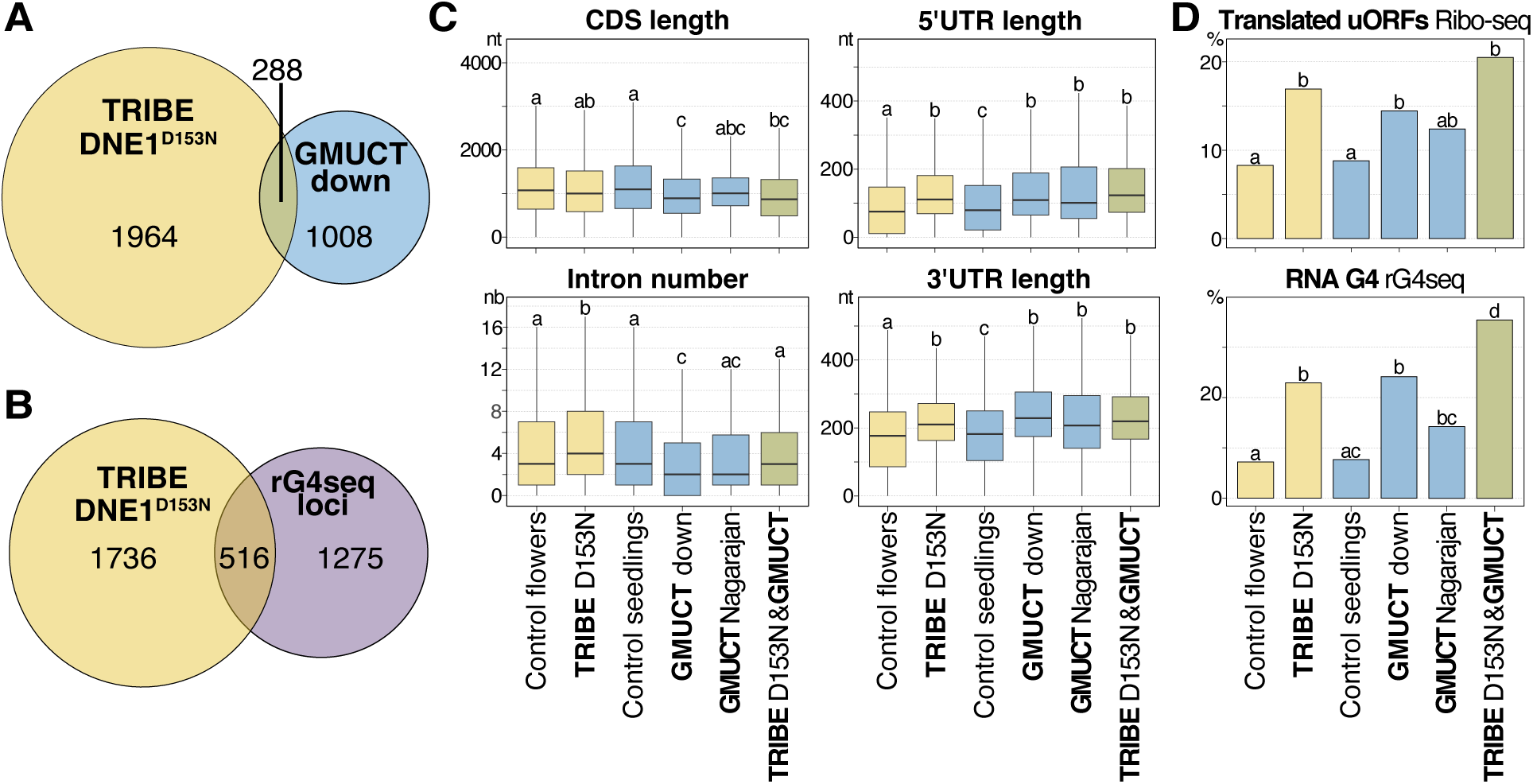
Analysis of mRNA features enriched in mRNAs identified in HyperTRIBE and GMUCT. (A) Venn diagram showing the overlap between loci edited by ADAR-DNE1^D153N^ and loci producing DNE1 dependent 5’P fragments. (B) Venn diagram showing the overlap between loci edited by ADAR-DNE1^D153N^ and transcripts containing validated RNA-G quadruplex (rG4). (C) Boxplot analysis of the number of introns and of mRNA, 5’ and 3’ UTR lengths for the DNE-dependent loci identified by the different methods. Significantly different values (adjpv < 0.001) are labelled by different letters (Wilcoxon rank sum test). D) Proportion of transcripts containing uORFs or rG4 in the different lists of DNE-dependent loci based on refs. Significantly different values (adjpv < 0.001) are labelled by different letters (two-samples z-test of proportions). In (C) and (D) the lists of transcripts expressed in flowers and seedlings are used as control.

### Mutations in *DNE1* and *DCP2* lead to synergistic transcriptomic changes

To better understand the impact and coordinated action of DNE1 and decapping on the transcriptome, we performed a transcriptomic analysis on a series of mutants including *dcp2* (*its1*, a previously described hypomorphic allele of *dcp2*), *dne1, dne1 dcp2* and *xrn4* (Fig. 4). Our working hypothesis from previous work and phenotypic analysis of these mutants predicts that combining mutations in *DNE1* and *DCP2* should synergistically affect the transcriptome and that *xrn4* and *dne1 dcp2* might affect some similar transcripts. Accordingly, we observed that whereas *dne1* and the weak allele of *dcp2* have a modest impact on the transcriptome (Fig. 4A), this effect is exacerbated in the two *dne1 dcp2* double mutant combinations (*dne1-2 dcp2* and *dne1-3 dcp2*, Fig. 4A, Supplemental Data Set S3). Overall, the most prominent trend observed in *dne1 dcp2* is upregulated transcripts and illustrate the synergistic effect of combining *dne1* and *dcp2* on the steady state level of specific mRNAs. Comparing these upregulated transcripts in *xrn4* and *dne1 dcp2*, two genetic backgrounds showing similar developmental defects, revealed that 51 transcripts were commonly deregulated in these mutants (Fig. 4B, 4C). These genes represent good candidates to identify genes involved in the phyllotactic defects observed. They notably include three bHLH transcription factors, PERICYCLE FACTOR TYPE-B 1 (PFB1: AT4G02590), LONESOME HIGHWAY LIKE 1 and 2 (LHL1: AT1G06150 and LHL2: AT2G31280). PFB1 is known to govern the competence of pericycle cells to initiate lateral root primordium, its involvement in organ emergence in the shoot is currently unknown (Zhang et al., 2021). LHL1 and LHL2 are known to regulate early xylem development downstream of auxin in roots and interestingly the use of an online tool to predict expression in the shoot apex indicate that both genes are expressed around the shoot apical meristem (Zhang et al., 2021); Supplemental Fig. S3). A fourth gene RAP2.4 for RELATED TO AP2 4 (AT1G78080) caught our attention. RAP2.4 it is an ethylene responsive factor, interestingly ERF12 another AP2 ethylene response factor was recently shown to be required for phyllotaxis (Chandler and Werr, 2020). These genes represent good candidates to better understand the importance of DNE1, DCP2 and XRN4 in phyllotaxis formation. Focusing on genes commonly upregulated in the two *dne1 dcp2* double mutants we asked whether some of them were identified as direct targets of DNE1 in either GMUCT or HyperTRIBE. We found that among these 68 genes 7 are found in GMUCT and 20 are found in HyperTRIBE for a total of 21 genes identified as putative direct targets of DNE1 including RAP2.4 identified in both approaches (Fig. 4D). This result highlights the redundancy of DNE1 and DCP2 in the regulation of gene expression and provides candidate genes to investigate the importance of these factors for phyllotaxis.

**Figure 4.**
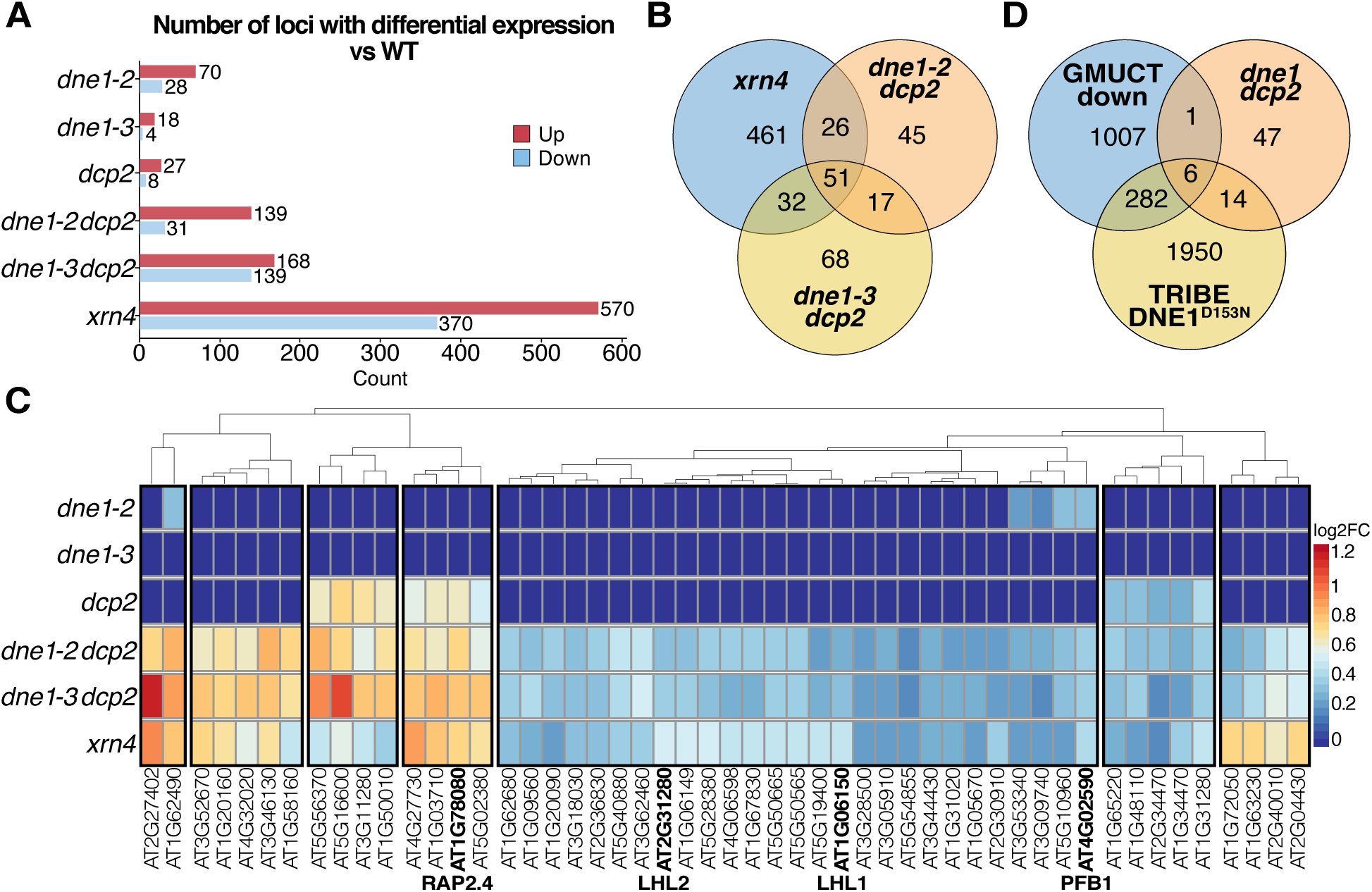
Transcriptomic analysis of *dcp2*, *dne1 dcp2* and *xrn4* mutants identify commonly deregulated transcripts. (A) Plot showing the number of differentially expressed genes in *dne1, dcp2, dne1 dcp2* and *xrn4* versus WT with adjPv<0.05 (n=3). (B) Venn diagram showing commonly upregulated loci between the two *dne1 dcp2* double mutants and *xrn4.* (C) Heatmap showing the mRNA accumulation pattern in *dne1, dcp2, dne1 dcp2* and *xrn4* for loci upregulated in both *dne1 dcp2* double mutants. (D) Venn diagram showing the overlap between upregulated loci in both *dne1 dcp2* double mutants and loci identified by GMUCT and HyperTRIBE.

### Differential sRNA populations can be instructive to identify targets of mRNA decay factors

Mutations in mRNA decay factors including *xrn4*, *dcp2* or *ski2* lead to the accumulation of 21 to 22 nt mRNA-derived siRNAs (Gregory et al., 2008; De Alba et al., 2015; Branscheid et al., 2015; Zhang et al., 2015). This phenomenon is due to the conversion of stabilized mRNA decay intermediates into siRNAs by the action of the RNA silencing machinery. Interestingly, several of these mRNA-derived siRNAs affect plant development as observed in *dcp2*, *xrn4 ski2*, *urt1 xrn4* (De Alba et al., 2015; Zhang et al., 2015; Scheer et al., 2021). Studying these siRNA populations have thus a double interest, it could help the identification of siRNAs potentially involved in the developmental defects appearing in corresponding mutants and it could allow the identification of mRNA targets of DNE1 and DCP2. To determine if the production of mRNA-derived siRNAs in RNA degradation mutants can be used as a criterion to identify targets of mRNA decay factors, we first analyzed small RNA populations accumulating in *xrn4* and *dcp2* (Fig. 5). XRN4 and DCP2 act sequentially in mRNA decay, the prediction is that they should accumulate populations of mRNA-derived siRNAs on similar loci. As expected, the main trend observed in *xrn4* and *dcp2* was upregulated mRNA-derived siRNAs populations (4737 loci in *xrn4*, and *2386* loci in *dcp2,* Fig. 5A, Supplemental Data Set S4). Interestingly, we observed a major overlap between siRNA loci in both mutants (with 2186 common loci, Fig. 5B). Of note, some of these loci are known *bona fide* targets of XRN4 including some of the first validated XRN4 targets, AT4G32020 and AT1G78080 (Souret et al., 2004). This first comparison shows that we can use mRNA-derived siRNA signatures differentially accumulating in RNA decay mutants to identify targets of mRNAs decay factors.

**Figure 5.**
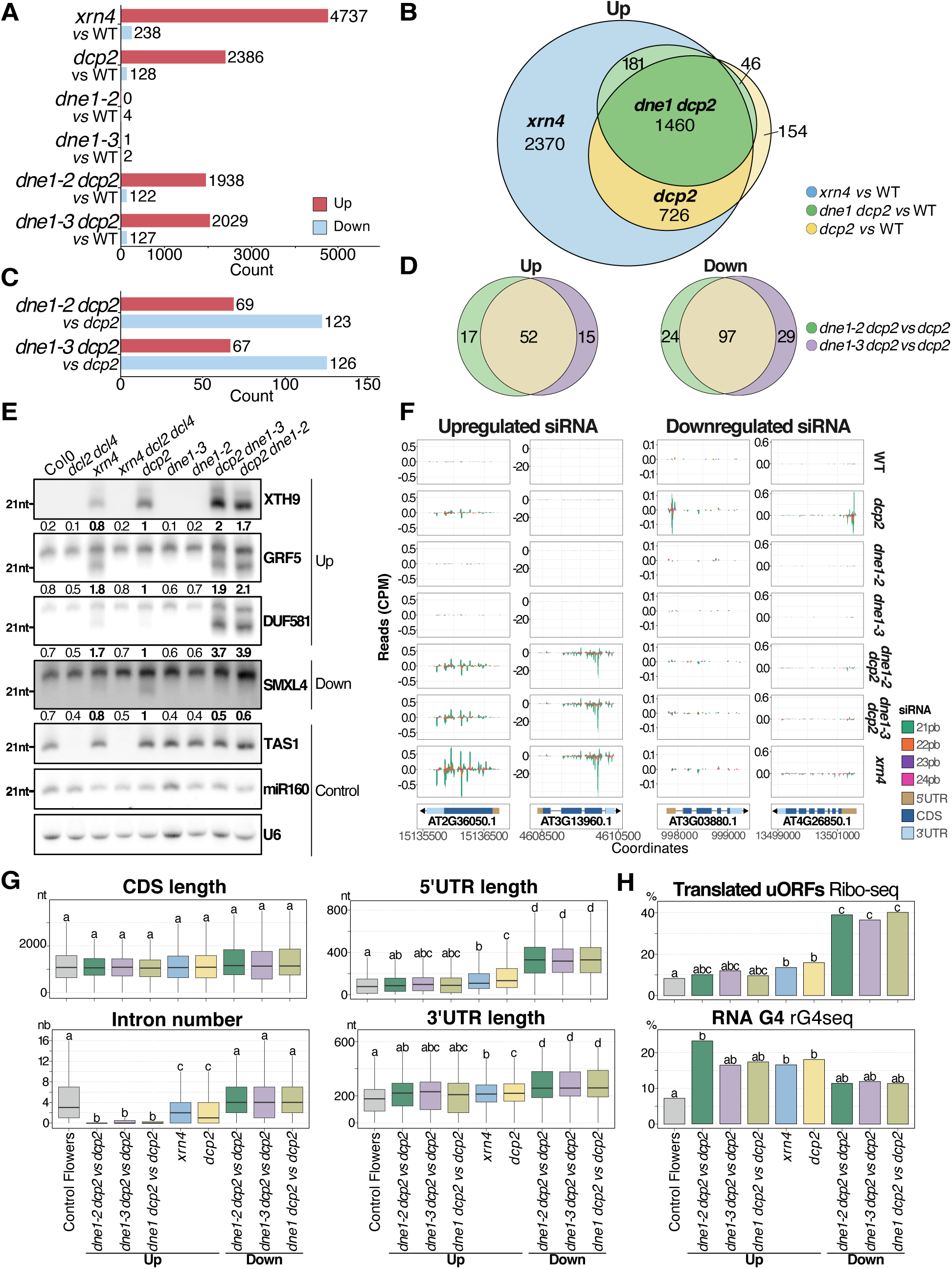
Differential analysis of small RNA accumulation in *dcp2*, *dne1 dcp2* and *xrn4* mutants. (A) Bar plots showing the output of the differential analysis of sRNA accumulation comparing mutants versus WT with adjPv<0.05 (n=3). (B) Venn diagram showing the overlap observed for upregulated sRNAs between different mutants. (C) Bar plots showing the output of the differential analysis of sRNA accumulation comparing *dne1 dcp2* versus *dcp2*. (D) Venn diagram showing the overlap observed for upregulated and downregulated sRNAs between the two *dne1 dcp2* double mutants. (E) Northern blot showing sRNA accumulation for loci differentially accumulating in *dne1 dcp2 vs dcp2.* The quantification is the mean and was performed with ImageJ on blots from three biological replicates. The 21nt size was determined by hybridization with an antisense probe targeting miR160. U6 was used as a loading control. (F) Plots showing the accumulation of mRNA-derived siRNAs along the transcripts for loci with upregulated and downregulated siRNAs. Datasets from the three biological replicates were pooled to generate these graphs. (G) Boxplot analysis of the number of introns and of mRNA, 5’ and 3’ UTR lengths for transcripts with differential sRNA accumulation in *xrn4*, *dcp2*, and *dne1 dcp2*. Significantly different values (adjpv < 0.001) are labelled by different letters (Wilcoxon rank sum test). (H) Proportion of transcripts containing uORFs or rG4 in the different lists of transcripts with differential sRNA accumulation. Significantly different values (adjpv < 0.001) are labelled by different letters (two-samples z-test of proportions). In (G) and (H) the list of transcripts expressed in flowers is used as control.

### Small RNA sequencing identifies DNE1-dependent small RNA populations

We used the same approach to identify mRNAs targeted by DNE1 by looking at mRNA-derived siRNA signatures differentially accumulating upon mutation of DNE1. In this analysis, we analyzed *dne1-2, dne1-3* and the corresponding *dne1 dcp2* double mutants. Globally, we found little changes in mRNA-derived siRNA accumulation in single *dne1* mutants and more changes in *dne1 dcp2* (Fig. 5A). This increase in the double mutant is largely due to the *dcp2* mutation as we observed a large overlap between sRNA populations upregulated in *xrn4*, *dcp2* and *dne1 dcp2* (Fig. 5B, 1460 loci).

This first analysis did not reveal a significant impact of mutation in DNE1 on siRNA accumulation. To investigate this point further we performed a differential analysis of siRNAs in *dne1 dcp2* using *dcp2* as a reference. In this analysis we identified two opposite trends, upregulated siRNA populations (69 loci in *dne1-2 dcp2* and 67 loci in *dne1-3 dcp2,* Fig. 5C) and downregulated siRNA populations (123 loci in *dne1-2 dcp2* and *126* loci in *dne1-3 dcp2*, Fig. 5C). An important overlap was observed between the two double mutants with 52 loci for upregulated siRNAs and 97 loci for downregulated siRNAs in common in both *dne1 dcp2* combinations (Fig. 5D). Both tendencies could be validated on a siRNA northern blot, which also illustrates that many of these siRNA species are produced in an *xrn4* mutant (Fig. 5E). Of note we analyzed in these blots triple *xrn4 dcl2 dcl4* mutants, which confirmed that theses siRNAs are produced by the RNA silencing machinery and involve the two main Dicer-like proteins involved in RNA silencing amplification DCL4 and DCL2. To better describe these patterns, we inspected the distribution of these siRNA on the transcripts. We observed that the siRNA distribution is different between upregulated and downregulated siRNAs. Upregulated siRNAs are mainly located on the CDS and 3’UTR (40/52, Fig. 5F Up, Supplemental Fig. S4A, Supplemental Data Set S5), in contrast downregulated sRNAs were mainly arising from 5’UTR (65/97; Fig. 5F Down, Supplemental Fig. S4 Down, Supplemental Dataset S5). We looked at the distribution of diverse mRNA features, including CDS, UTR length and intron number, in the loci associated with each trend compared to overall expressed genes (Fig. 5G). The most striking feature for loci with upregulated siRNAs was a strikingly low intron number, identifying those loci as intron-poor mRNAs. In contrast loci with less siRNAs possess the same number of introns than other expressed transcripts and had slightly longer 3’UTR and strikingly longer 5’UTR. We then tested whether loci with differential siRNA patterns were particularly enriched in transcripts containing translated uORFs in 5’UTR or rG4, as observed in mRNA identified as DNE1 targets in GMUCT and HyperTRIBE. The most striking result of this analysis appeared for loci with downregulated siRNAs in *dne1 dcp2* versus *dcp2* (already identified to harbor dramatically longer 5’UTR), which were noticeably enriched in mRNA containing translated uORFs (Fig. 5H). In terms of siRNA accumulation, the general trend for upregulated and downregulated siRNAs is the exacerbation or attenuation of siRNA populations observed in *dcp2* (Fig. 5F), suggesting that both DCP2 and DNE1 target those transcripts. Despite the relatively low number of differential loci found in siRNA-seq we found an overlap between loci found in siRNA sequencing, HyperTRIBE and GMUCT (Fig. 6). Overall, 44 loci showing differential siRNA patterns were identified as DNE1 targets by GMUCT or HyperTRIBE suggesting that they represent *bona fide* DNE1 targets. One of the most striking examples of this trend is the RAP2.4 gene AT1G78080, which was recovered in every HTS methods, it is heavily edited by DNE1-D153N mainly in the CDS (Supplemental Fig. S1), it presents both upregulated and downregulated RNA fragments in *dne1 xrn4* in GMUCT (Supplemental Fig. S2B), its mRNA is upregulated in *dne1 dcp2* in RNA-seq (Fig. 4C) and it produces less siRNA in *dne1 dcp2 versus dcp2* in its 5’UTR (Supplemental Fig. S4B). These observations indicate that every method used in this study, despite the fact that they monitor completely different features, has the potential to identify DNE1 targets highlighting the added value of our multi-transcriptomic approach.

**Figure 6.**
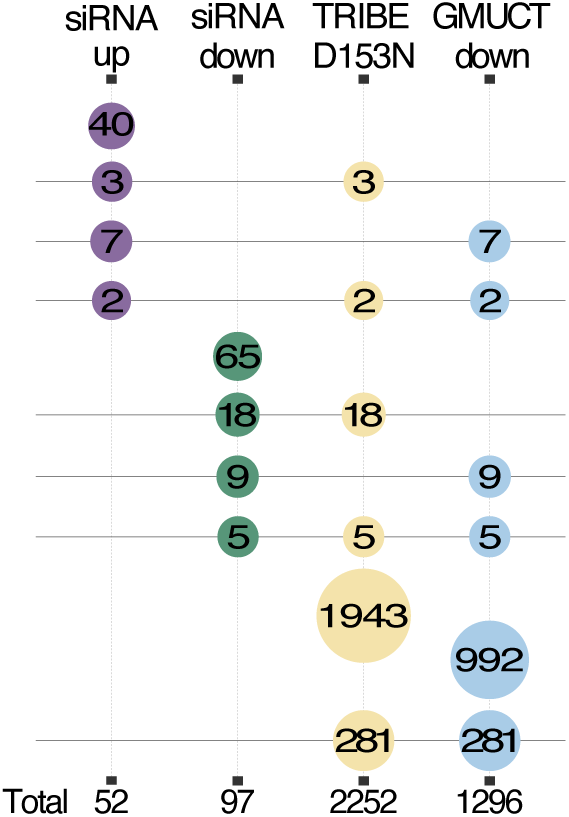
Diverse HTS techniques identifiy specific and common mRNAs influenced by DNE1. Bubble chart showing the extent of intersection between the list of loci identified by sRNA-seq, HyperTRIBE and GMUCT. Each column corresponds to a list of loci and each row correspond to a possible intersection. Bubbles indicate the number of loci for each intersection with colors showing the number of related lists.

## Discussion

In this work we combined *in vivo* RNA editing by HyperTRIBE and RNA degradome sequencing by GMUCT to identify targets of the endoribonuclease DNE1. The advantage of HyperTRIBE is to identify mRNAs contacting DNE1 but its intrinsic limitation is that is does not give any indication regarding mRNA cleavage by DNE1. The advantage of using GMUCT is to identify mRNAs cleaved by DNE1 but its limitation is that this identification is only possible if the corresponding RNA degradation products are sufficiently stable. These limitations are solved when combining HyperTRIBE with GMUCT giving access to independent lists of targets. In addition, the overlap between the two methods identifies a refined list of mRNAs contacting and cleaved by DNE1.

In our work we also interrogated the influence of DNE1 and DCP2 on mRNA fate using transcriptomics and small RNA deep sequencing in the *dne1 dcp2* double mutant. While transcriptomics identified mRNAs with altered steady state levels in *dne1 dcp2*, the most interesting information regarding DNE1 action and coordination with DCP2 came from the study of mRNA-derived siRNAs. The identification of differential mRNA-derived siRNAs in *dne1 dcp2* compared to *dcp2* supported the hypothesis of their action on similar transcripts. We consider changes in mRNA-derived siRNA production in *dne1 dcp2* as a readout of changes in mRNA fate when DNE1 function is abrogated. Unexpectedly, two trends appeared in this analysis, upregulated siRNAs and downregulated siRNAs. We propose a model to explain the appearance of these two opposite trends. Our interpretation of this result is that both trends appear on mRNAs targeted by DNE1. This is coherent with the presence of some of these loci in GMUCT and/or HyperTRIBE. Upregulated siRNAs are produced all along the transcripts in *dcp2*, suggesting that they are produced from full-length mRNAs that are stabilized when DCP2 function is affected. In *dcp2*, DNE1 cleaves a pool of these transcripts reducing the pool of full-length transcripts available for decapping. When DNE1 is mutated the pool of full-length transcripts increases leading to increased targeting by DCP2. This increased targeting by DCP2 leads in *dne1 dcp2* to increased proportion of stabilized full-length mRNAs and increased siRNA accumulation, likely produced from full-length capped mRNAs (Fig. 7, panel A).

**Figure 7.**
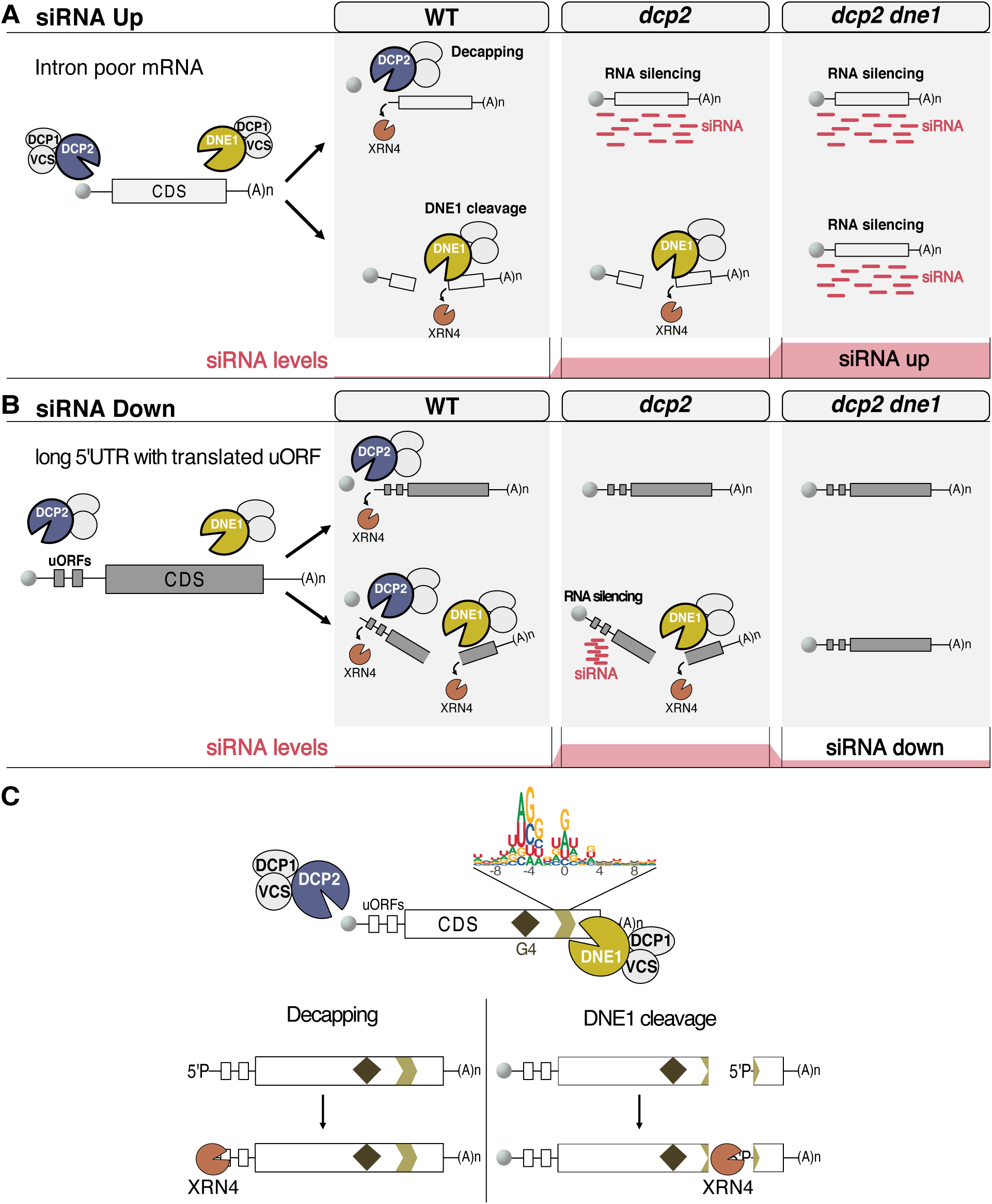
Models of DNE1 and DCP2 coordinated action on mRNAs. (A), (B) Integrated models for the action of DNE1 and DCP2 on mRNA-derived siRNAs production. (C) Integrated model built from the HyperTRIBE and GMUCT data. The model shows interaction and action of DNE1 in the CDS on sites with preferred nucleotide composition. Enriched features in DNE1 targets including RNA-G4 and translated uORFs are depicted.

In contrast downregulated siRNAs are mainly produced in discrete positions from 5’UTRs. Our interpretation is that they are not produced from full-length mRNA but from stabilized DNE1 cleavage products. In this case abrogating DNE1 action in *dne1 dcp2* leads to the reduction in the accumulation of DNE1 cleavage products and a reduction in mRNA-derived siRNA production from these products (Fig. 7B). This interpretation implies that DNE1 cleavage products can be decapped by DCP2. In addition of this mechanistic model, we found that these two lists of mRNAs are enriched for very different features. mRNAs with upregulated siRNAs are strikingly intron-poor mRNAs. This is reminiscent of previous studies on transgenes, in which it was described that introns protect transgenes from RNA silencing activation (Christie et al., 2011). We propose that these mRNAs are specifically prone to siRNA production due to their low introns number, this low intron number trend was also identified but to a slightly lower extent for mRNAs producing siRNAs in *xrn4* and *dcp2* in our data (Fig. 5G). This results strongly support the hypothesis that in RNA degradation mutants, introns protect mRNAs from RNA silencing activation as previously observed in WT plants (Christie et al., 2011). In contrast mRNAs with downregulated siRNAs had similar intron numbers as overall expressed mRNAs but were characterized by strikingly longer 5’UTR. Interestingly, we found that these long 5’UTR were significantly enriched in translated uORFs, coinciding with the sites of siRNA production in *dcp2.* We can speculate that the translation of these uORFs might further stabilize these cleavage products allowing them to partially escape 3’ to 5’ degradation leaving enough time for them to be detected and processed by the RNA silencing machinery leading to siRNA production.

How DNE1 recognizes its targets and what is the trigger to induce DNE1 mediated RNA degradation are fundamental questions to be addressed in future studies. Definitive answers to these questions will require more work but the identification of enriched features among DNE1 targets can be instructive to formulate hypothesis. First, we identified that transcripts identified in the HyperTRIBE and GMUCT approaches are enriched in translated uORFs and rG4. Remarkably, this trend is exacerbated in the highest confidence DNE1 targets commonly identified in GMUCT and HyperTRIBE (Fig. 3C). This observation suggests that translated uORFs and rG4 might promote targeting and cleavage by DNE1. Of note, for many transcripts identified to contact DNE1 in HyperTRIBE, we did not detect differential RNA fragments in GMUCT. A possible explanation for this discrepancy is that DNE1 might contact both targets and non-targets in a scanning mode, looking for cleavage inducing features. Hallmarks of this potential scanning can be found in the HyperTRIBE results (Supplemental Fig. S1) as some targets were edited all along the CDS. Translated uORFs are known to regulate gene expression by impairing translation of the main ORF. In this scenario, DNE1 would scan mRNAs containing translated uORF with inefficient translation of the main ORF. The inefficient translation of the main ORF could allow the formation of tertiary structures in the main ORF including rG4. While DNE1 scans these mRNAs it encounters rG4 or other structures, they are recognized by the OST-HTH domains of DNE1, identified as G rich and rG4 interacting domains *in vitro* and induce cleavage by DNE1. Our analysis of DNE1 cleavage sites revealed a biased nucleotide composition. The identification of this nucleotide preference at DNE1 cleavage site is fundamentally different from the previous identification of an enriched G-rich motif (YGGWG) in the vicinity of DNE1 cleavage site (Nagarajan et al., 2023). While the YGGWG motifs are found at various positions surrounding the cleavage site, the nucleotide preference identified here occurs at very precise position on and around cleavage sites. Interestingly, a similar nucleotide preference appeared when we performed the logo analysis on the 224 DNE1 targets identified in the previous study (Supplemental Fig. S2C), validating the efficiency of our identification of 1295 DNE1 target in GMUCT and the relevance of this logo. This observation reveals that DNE1 does not cleave mRNAs at random sequences and support the hypothesis that DNE1 have nucleotide context preferences for its endonuclease activity. To sum up the previous observations we build a final model illustrating the coordinated action of DNE1 and DCP2 in the degradation of DNE1 targets (Fig 7C).

Overall, our study greatly increases the spectrum of potential DNE1 targets. It will be crucial to pursue the efforts and to start investigating how DNE1 regulates specific processes at the tissues level. We previously showed that together with DCP2, DNE1 is required for phyllotaxis, the formation of precise developmental patterns at the shoot apex. Our current work provides a first extended list of DNE1 targets that can be searched to identify novel regulators of phyllotaxis. Which of these targets are locally expressed in developing primordia? How their expression is altered upon mutation in DNE1 and DCP2 and what are the physiological changes in the shoot apex in *dne1 dcp2*? Answers to these questions will be crucial to better understand the importance of these factors for phyllotaxis and combining the study of *dne1 dcp2* and *xrn4* will reveal the overall importance of RNA degradation in the control of the homeostasis of key regulators of phyllotaxis.

## Materials and methods

### Plant materials and growth conditions

*Arabidopsis thaliana* mutants and WT lines were in the Columbia-0 (Col-0) ecotype. Mutants used in this study were all previously described: *dne1-1* (Salk_132521); *xrn4-3* (SALK_014209); *dcl2-1* (SALK_064627), *dcl4-2* (GABI_160G05), *dne1-2* and *dne1-3* were produced by the CRISPR/Cas9 system (Schiaffini et al., 2022). Transgenic lines produced in the HyperTRIBE strategy were in the *dne1-3* mutant background. The plant material used for RNA-seq, small RNA-seq and HyperTRIBE were grown on soil in 16/8h light/dark conditions until flowering and unopened flower buds were collected. The plant material used for GMUCT were seedlings grown on Murashige and Skoog (MS) medium (MS0255 Duchefa, 0,7% w/v agar, pH 5.7). Seeds were sterilized with bleach/ethanol solution (0,48% / 70%) on shaker for 10min, and then wash with 70% ethanol. The seed were rinse twice with sterile water. After 24h of stratification at 4°C seedlings were grown in 16/8 h light/dark conditions at 21°C for 10-d and transferred into liquid half-strength MS medium. The seedlings were collected for RNA extraction after incubation at 40 rpm under constant light for 24h.

### Constructs produced for HyperTRIBE

p35S:FLAG-ADARcd^E488Q^-DNE1-35ST (F-ADAR-DNE1), p35S:FLAG-ADARcd^E488Q^-DNE1^D153N^-35ST (F-ADAR-DNE1^D153N^), ADARcd^E488Q^ (p35S:FLAG-ADARcd^E488Q^-35ST (FLAG-ADAR). Constructs were produced by overlap-extension PCR (Bryksin and Matsumura, 2013) to fuse the ADAR sequence to DNE1 followed by Gateway® recombination in pH2GW7. All final constructs were verified by Sanger sequencing and mobilized into *Agrobacterium tumefaciens* (GV3101 pMP90) chemically competent cells. Transgenic lines were generated by floral dip (Clough and Bent, 1998) of *dne1-3* with *A. tumefaciens* GV3101 bearing pH2GW7 F-ADAR-DNE1, F-ADAR-DNE1^D153N^ and FLAG-ADAR. Selection of primary transformants (T1) was done by hygromycin to select five independent lines for each type of transgene. Expression levels were assessed by western blot using anti-FLAG M2 antibodies. (Primers used in the study present supplemental table S1)

### Total RNA extraction

Total RNA was extracted using Tri-Reagent (Molecular Research Center, Inc., Cincinnati, OH, USA) according to the manufacturer’s instructions, followed by acidic phenol chloroform extraction and RNA precipitation with ethanol. The samples were then treated with DNase I (Thermo Fisher Scientific) according to the manufacturer’s instructions.

### RNA degradome library preparation

Poly(A)+ RNA isolated from 11 days old whole seedlings were used to generate GMUCT libraries according to the published protocol (Carpentier et al., 2021). Libraries were sequenced on Illumina HiSeq 2500 in a 50 nt single-end mode.

### Computational analysis of RNA degradome data

GMUCT libraries were aligned to TAIR10 genome with hisat2. The coverage of 5’ reads position (for both strands) were extracted using bedtools genomecov from the bam files. A differential expression analysis was performed between *xrn4* and *xrn4 dne1* (3 replicates per sample) using the DEXSeq R package with the following design: ~ sample + base + condition:base. All the scripts are available at https://github.com/ibmp/dne1_2024.

### HyperTRIBE library preparation

The HyperTRIBE analysis was performed on five independent lines of F-ADARcd^E488Q^ (control), F-ADARcd^E488Q^-DNE1 and F-ADARcd^E488Q^-DNE1^D153N^ used as five biological replicates. Purified total RNAs were quantified by Qubit (Invitrogen) fluorimeter, quality was assessed using Bioanalyzer 2100 (Agilent) system. Six hundred nanograms of RNAs were used for library preparation with the TruSeq® Stranded mRNA Library Prep following manufacturer’s instructions. Libraries were sequenced by paired-End (2×100bases) on an Illumina HiSeq 4000. Sequencing was performed by the GenomEast platform.

### Computational analysis of HyperTRIBE

Sequencing data were aligned to the TAIR10 reference genome with hisat2 using the following options:“-t -k 50 --max-intronlen 2000 --rna-strandness RF --no-unal”. The analysis was conducted following the steps described here https://github.com/sarah-ku/hyperTRIBER. In short, the bam files were split by strand and a single mpileup file was generated from all the files with samtools. The mpileup file was then converted using the RNAeditR_mpileup2bases.pl script. The resulting output was further analyzed in R with the hyperTRIBER package. Only A-to-G edits were selected.

### RNAseq library preparation

The RNAseq analysis was performed on biological triplicates of inflorescence of the WT, *its1* (*dcp2*), *dne1-2*, *dne1-3*, *xrn4-3* and two double mutant *its1 dne1-2* and *its1 dne1-3*. Purified total RNAs were quantified by Qubit (Invitrogen), RNA quality was tested using Bioanalyzer 2100 (Agilent) system. Six hundred nanograms of RNAs were used for library preparation with the TruSeq® Stranded mRNA Library Prep using manufacturer’s instructions. Libraries were sequenced by single read (1×50bases) with an Illumina HiSeq 4000. Sequencing was performed by the GenomEast platform.

### Computational analysis of RNAseq

Reads were first aligned to the TAIR10 reference genome using hisat2 aligner with the following options:

--max-intronlen 2000 -q --rna-strandness R --passthrough --read-lengths 50

Then, read counts were extracted for each representative transcript using FeatureCounts and a differential expression analysis was performed in R with the DESeq2 package. For all analyses, we used the most representative gene isoform (described in the TAIR10_representative_gene_models file).

### sRNAseq library preparation

Transcriptomic analysis was performed on biological triplicates of inflorescence of the wild type (col-0), *its1* (*dcp2*), *dne1-2*, *dne1-3*, *xrn4-3* and two double mutant *its1 dne1-2* and *its1 dne1-3*. Purified total RNAs were quantified by Qubit (Invitrogen) fluorimeter, RNA’s quality was tested using Bioanalyzer 2100 (Agilent) system. Six hundred nanograms of RNAs were used for libraries preparation with the NEBNext® Multiplex Small RNA Library Prep Set for Illumina® using manufacturer’s instructions. Libraries were sequenced by single read (1×50bases) with an Illumina HiSeq 4000. Sequencing was performed by the GenomEast platform.

### Computational analysis of sRNAseq

Raw reads were trimmed using trimgalore with the following options: “-q 30 --max_n 5 --max_length 30”. The resulting clean reads were mapped to TAIR10 reference genome with the following options: “-v 1 --best --strata -k 10”. The sRNA counts per size on each TAIR10 representative transcripts were extracted from each bamfile with ShortStack using the following options: “--nohp --dicermin 15 --dicermax 30”. To study mRNA-derived siRNAs, a differential expression analysis was done with DESeq2 using as counts the sum of 21 and 22nt long sRNAs in each transcript features. Extraction of Dicercall 21-dependent transcripts: the bam files from all replicates (3 replicates per sample) were merged into a single bam per sample. ShortStack was run on each merged bam. Loci identified as “DicerCall21” by ShortStack were extracted from the results. Subsequently, we selected loci that were found in at least 3 conditions out of 7 as DicerCall 21-dependent transcripts, resulting in a list of 7935 AGI.

### Low molecular weight northern blot

For this analysis we used 40ug of total RNA resuspended in sRNA loading buffer (4X: 50% glycerol, 50mM Tris pH 7.7, 5mM EDTA, 0.03% bromophenol Blue). The RNA was denatured at 95°C for 5min prior to loading in a prewarmed 17.5% acrylamide:bis 19:1; 7M urea, 0.5X TBE gel, electrophoresis was performed in 0.5 TBE at 80V for 5h. RNA was transferred onto an Amersham Hybond-NX membrane at 300mA in 0.5x TBE for 1h at 4°C. The membrane was chemically crosslinked with EDC (1-ethyl-3-(3-dimethylaminopropyl) carbodiimide) for 1h30 at 60°C. After crosslinking, the membrane was rinsed with water and incubated at 42°C for 45min in PerfectHyb^TM^ plus hybridization buffer. For probes produced by random priming, the purified PCR products were radiolabeled using the Prime-a-Gene® Labeling System according to the manufacturer’s instructions. For probes produced by end labeling, the primers were radiolabeled using the Thermo Scientific™ T4 Polynucleotide Kinase according to the manufacturer’s instructions. Radiolabelled probes were added directly in the buffer and the membrane was incubated overnight (O/N) with the probe at 42°C. The membrane was washed with 2xSSC (0.3M NaCl, 30mM sodium citrate) 2% SDS three times 20 min at 50°C. Signal intensities were analyzed using the Typhoon system (GE Health Sciences). Membranes were stripped in boiling 0.1% SDS three times 20min. Northern blot results presented are representative of 3 biological replicates. Primers used for probe preparation are listed in supplemental table S1.

### Protein extraction and Western blotting

Total protein was extracted using Tri-Reagent (MRC). Five flower buds were ground in 300 µl TRI-Reagent. After mixing 60 µl of chloroform were added then the sample is incubated 15 min at room temperature then centrifugated 15 min. After removing the aqueous phase, DNA is precipitated by adding 100µl ethanol, incubating for 15min and centrifuging for 15min at 18,000g. The supernatant was then recovered, and the proteins were precipitated by adding 3V of 100% acetone, followed by 5min incubation on ice. After centrifugation 1min at 5000g, the pellet was washed once with 80% acetone. The pellet was then recovered in SDS-urea buffer. (62.5 mM Tris pH 6.8, 4 M urea, 3% SDS, 10% glycerol, 0.01% bromophenol blue). The samples were separated by SDS-PAGE and transferred to a 0.45 μm Immobilon-P PVDF membrane (Millipore). The membrane was incubated 2h at 4°C with ANTI-FLAG antibodies® M2-peroxydase (Sigma-Aldrich, used at 1/ 1000 dilution). The antibodies were detected by using Lumi-Light Western Blotting Substrate (Roche). Pictures were taken with a Fusion FX camera system (Vilber). The PVDF membranes were stained with 0.1% Coomassie Brilliant Blue R-250, 9% acetic acid, 45.5% ethanol) to monitor loading.

### Comparison of HTS datasets with transcript characteristics

The number of introns and the length of CDS and UTRs used for the comparison were based on the TAIR10 annotation for representative transcripts. The proportion of mRNA containing uORFs and rG4 were retrieved from Hu et al. 2016 and Yang et al. 2020, respectively. For the control lists, we used the lists of transcripts detected by RNAseq in WT flowers (this paper, Supplemental Data Set S3) and in WT seedlings (Schiaffini et al. 2022). Boxplots shown Fig.3 and 5 displays the median, first and third quartiles (lower and upper hinges), the largest value within 1.5 times the interquartile range above the upper hinge (upper whisker) and the smallest value within 1.5 times the interquartile range below the lower hinge (lower whiskers). In Fig.3C and 4C, statistical analysis was performed using Pairwise Wilcoxon Rank Sum Tests with data considered as unpaired (non-parametric test, two-tailed). In Fig.3D and 4D, a two-samples z-test of proportions was applied. For all statistical analysis, an adjusted p-value (fdr) of 0.001 was defined as threshold of significance. Plots and statistics were performed using R (v4.2.2), and R packages ggplot2 (v3.4.5) and stats (v4.2.2). Scripts are available in Github (https://github.com/hzuber67/Feature_analysisDNE1).

## Accession numbers

Raw and processed sequences of RNAseq, SmallRNAseq, HyperTRIBEseq, and GMUCT libraries (Supplemental Data Set S1 to S4) are available at the National Center for Biotechnology Information (NCBI)-Sequence Read Archive (SRA) under the accession number PRJNA995202. Sequence corresponding to genes mentioned in this article can be found in the Arabidopsis Information Resource (TAIR _ https://www.arabidopsis.org/) under the following accession numbers: AT2G15560 (DNE1); AT4G03210 (XTH9); AT3G13960 (GRF5); AT5G20700 (DUF581); AT4G29920 (SMXL4); AT1G54490 (XRN4); AT3G03300 (DCL2); AT5G20320 (DCL4); AT5G13570 (DCP2); AT1G06150 (LHL1); AT2G31280 (LHL2/LL2); AT1G78080 (RAP2-4). CG12598 NM_001297862 (ADAR isoform N).

## Acknowledgements

The authors thank Peter Brodersen and Laura Arribas Hernandez for advices and for sending the plasmids containing the ADARcd domain used in HyperTRIBE and Sarah Rennie for support in using the HyperTRIBER pipeline. Sandrine Koechler from the Plateforme d’analyse d’expression génique for help in RNA-seq libraries preparation.

## Author contribution

**Damien Garcia**: research design, conceptualization, data interpretation, initial analysis of HTS data, writing – original draft, writing – review and editing. **Aude Pouclet**: performed the experiments, collected and interpreted the data, writing – review and editing. **David Pflieger**: analysis and visualization of HTS data, writing – review and editing. **Rémy Merret:** performed the GMUCT, writing – review and editing; **Marie-Christine Carpentier:** performed the primary analysis of GMUCT data, writing – review and editing. **Marlene Schiaffini** produced mutant combinations, writing – review and editing. **Hélène Zuber** performed statistical analysis and comparison of HTS datasets, writing – review and editing. **Dominique Gagliardi:** Conceptualization, data interpretation, writing – review and editing.

## Supplemental data

**Supplemental Figure S1.** Schemes showing the editions by ADAR-DNE1^D153N^ on 11 transcripts illustrating the preferential edition in the CDS.

**Supplemental Figure S2.** Profiles of 5’P fragments accumulation in GMUCT on representative examples. (A) Plots showing the repartition of 5’P on loci presenting only downregulated 5’P fragments. (B) Plots showing the repartition of 5’P on five loci presenting both downregulated and upregulated 5’P fragments. (C) Logo analysis performed on the 224 DNE1 targets identified in Nagarajan et al 2023.

**Supplemental Figure S3.** Predicted expression patterns of AT2G31280 (LL2) and AT1G06150 (LHL1) in the shoot meristem of *Arabidopsis thaliana* using the 3D flower meristem tool from single cell experiments performed in Neumann et al 2022.

**Supplemental Figure S4.** Representative examples of transcripts showing differential accumulation of mRNA-derived siRNAs between *dcp2* and *dne1 dcp2*. (A) Plots showing the accumulation of mRNA-derived siRNAs along the transcripts for upregulated siRNAs. (B) Plots showing the accumulation of mRNA-derived siRNAs along the transcripts for downregulated siRNAs.

**Supplemental Yable S1.** Primer list.

**Supplemental Data Set S1.** HyperTRIBE data.

**Supplemental Data Set S2.** GMUCT data.

**Supplemental Data Set S3.** RNAseq data.

**Supplemental Data Set S4.** sRNAseq data.

**Supplemental Data Set S5.** Localization of differential sRNA on transcripts showing differential accumulation in *dne1 dcp2 vs dcp2*.

**Supplemental Data Set S6.** Statistics for feature enrichment analysis in Figure 3 and 5.

**Supplemental Data Set S7.** Lists of loci used to identify mRNA features in Figure 3 and 5.

